# Comparison of human and murine enteroendocrine cells by transcriptomic and peptidomic profiling

**DOI:** 10.1101/374579

**Authors:** Geoffrey P Roberts, Pierre Larraufie, Paul Richards, Richard G Kay, Sam G Galvin, Emily L Miedzybrodzka, Andrew Leiter, H. Joyce Li, Leslie L Glass, Marcella KL Ma, Brian Lam, Giles SH Yeo, Raphaël Scharfmann, Davide Chiarugi, Richard H Hardwick, Frank Reimann, Fiona M Gribble

**Affiliations:** Wellcome Trust – MRC Institute of Metabolic Science, University of Cambridge, UK; Cambridge Oesophago-gastric Centre, Addenbrooke’s Hospital, Cambridge, UK; INSERM U1016, Institut Cochin, Université Paris-Descartes; Division of Gastroenterology, Department of Medicine, University of Massachusetts Medical School, Worcester, Massachusetts, United States

**Keywords:** Gut hormones, GLP-1, enteroendocrine, PYY, GIP, bariatric surgery, transcriptome, peptidome, LC-MS, RNAseq

## Abstract

Enteroendocrine cells (EECs) produce hormones that regulate food absorption, insulin secretion and appetite. Both EECs and their peptide products are foci of drug discovery programmes for diabetes and obesity. We compared the human and mouse EEC transcriptome and peptidome to validate mouse as a model of the human enteroendocrine axis. We present the first RNA sequencing analysis of human EECs, and demonstrate strong correlation with mouse, although with outliers including some low abundance G-protein coupled receptors. Liquid chromatography mass spectrometry (LC-MS) identified peptide hormone gradients along the human and mouse gut that should enhance progress in gut physiology and therapeutics.

## Introduction

Enteroendocrine cells (EEC) are specialised hormone secreting cells in the intestinal epithelium which monitor the quality and quantity of ingested foods. They produce at least 20 different hormones, mostly peptides, that act in concert to coordinate digestion, peripheral nutrient disposal and appetite through actions at local and distant target tissues. In the field of human metabolism, glucagon-like peptide-1 (GLP-1) and peptideYY (PYY) have raised particular interest because of their central and pancreatic actions controlling food intake and insulin secretion. GLP-1 based drugs are widely used for the treatment of diabetes and obesity, and new gut hormone based therapeutics are under development, aiming to mimic the unrivalled effectiveness of gastric bypass surgery on weight loss and type 2 diabetes resolution(2).

Our understanding of the physiology and pharmacology of human EECs is currently limited by a lack of methods to identify and characterise this scattered cell population which only comprises ^~^1% of the intestinal epithelium(3). By contrast, recent years have witnessed substantial progress in our understanding of murine EEC physiology, facilitated by the generation of transgenic mice with fluorescently labelled EECs that enable cell identification and functional characterisation through a range of approaches including fluorescence-activated cell sorting (FACS), transcriptomics and live cell imaging(3-8). The lack of an evidence-based platform for predicting the translatability of these findings from mouse to human represents, however, a significant obstacle in drug discovery programmes. Whilst G-protein coupled receptors (GPCRs) that have been identified and characterised in murine EECs represent promising candidates for therapeutic approaches to enhance endogenous gut hormone secretion, early validation in a human model is strongly advisable(9).

Our objective was to compare mouse and human EECs at the transcriptomic and peptidomic levels. For transcriptomics, we purified and RNA sequenced two subsets of human EECs distinguished by their production (or not) of GLP-1, and compared the results with matching murine cell populations purified from transgenic mouse models. Sequencing depth in both species was sufficient to enable the identification of low abundance transcripts such as GPCRs and ion channels. We optimised protocols for peptide extraction and liquid chromatography / tandem mass spectrometric (LC-MS/MS) analysis of mucosal tissue homogenates, and identified exact sequences of the different gut peptides produced, and their relative quantities along the length of the GI tract in human and mouse.

## Results

### Cell and RNA collection for transcriptomics

Tissue was obtained from 11 human participants recruited from sites in Cambridge and Paris (Supplementary table 1). Jejunum samples were collected from all participants, and paired ileum and jejunum were obtained from 2 transplant donors. Tissue pieces were digested, paraformaldehyde (PFA)-fixed and stained for Chromogranin A (CHGA) and Secretogranin 2 (SCG2) as general markers for EECs, and for GLP-1 as a marker of the EEC-subpopulation known as L-cells. By flow cytometry (Supplementary Figure 1A-C), we collected pooled cell populations that were (i) positive for CHGA, SCG2 and GLP-1 (henceforth named GCG+), (ii) positive for CHGA and SCG2 but negative for GLP-1 (henceforth named GCG-), and (iii) negative for all 3 markers (i.e. non-endocrine lineage cells, henceforth named negative). The GCG+ (L-cell) population represented ^~^0.2% of all single cells examined, and the ratio of GCG+ to GCG-cells was ^~^1:5.

For the cross-species comparison, we collected unfixed murine EEC populations from the upper small intestine of the mouse strain GLU-Venus (n=3) to identify *Gcg*-expressing L-cells (Supplementary Figure 1E-G)(8), and of NeuroD1-Cre/EYFP mice(7) (n=3) to identify the total EEC population (Supplementary Figure 1I-K). NeuroD1 is a well characterised EEC transcription factor found in the same cells as *Chga* in a previous single cell analysis of murine EECs(10). GLU-Venus positive cells represented ^~^0.2% of singlets, and NeuroD1 positive cells ^~^0.6% of singlets.

RNA extracted from the purified fixed human cell populations had RIN values of only 2-3, with most RNA fragments being 25-500 bases in length (Supplementary Figure 1D), but we hypothesised that the short fragments might be sufficient for RNA sequencing using random primers. After RNA sequencing there were high proportions of unmatched reads (up to 50% in some samples), likely due to the PFA modified input RNA, but in all cases sequencing was achieved to a depth of at least 8.5×10^5^ matched reads per sample. RNAs from murine cell populations purified without fixation had RIN values of >7 (Supplementary Figure 1H,L) and were sequenced to >5×10^6^ aligned reads per sample.

### Transcriptomics of human EEC populations

In total we obtained individual RNA sequencing data from GCG+, GCG- and negative cells from each of 11 human jejunum samples and 2 human ileum samples. Principle component analysis (PCA) of the top 500 differentially expressed genes across all samples showed separation of the EECs (GCG+ and GCG-) from negative cells on the first component, and of GCG+ cells from GCG-cells on the second component (Figure 1A). Pairwise analysis of key genes differentiating the cell populations from jejunum was performed using a DESEQ2 model, and the normalised results for the 500 most differentially expressed genes are presented in a heatmap (Figure 1B) which displays clear transcriptomic differences between the GCG+, GCG- and negative cells(11). PCA did not demonstrate clustering of samples by BMI of the donor.

**Figure 1.**
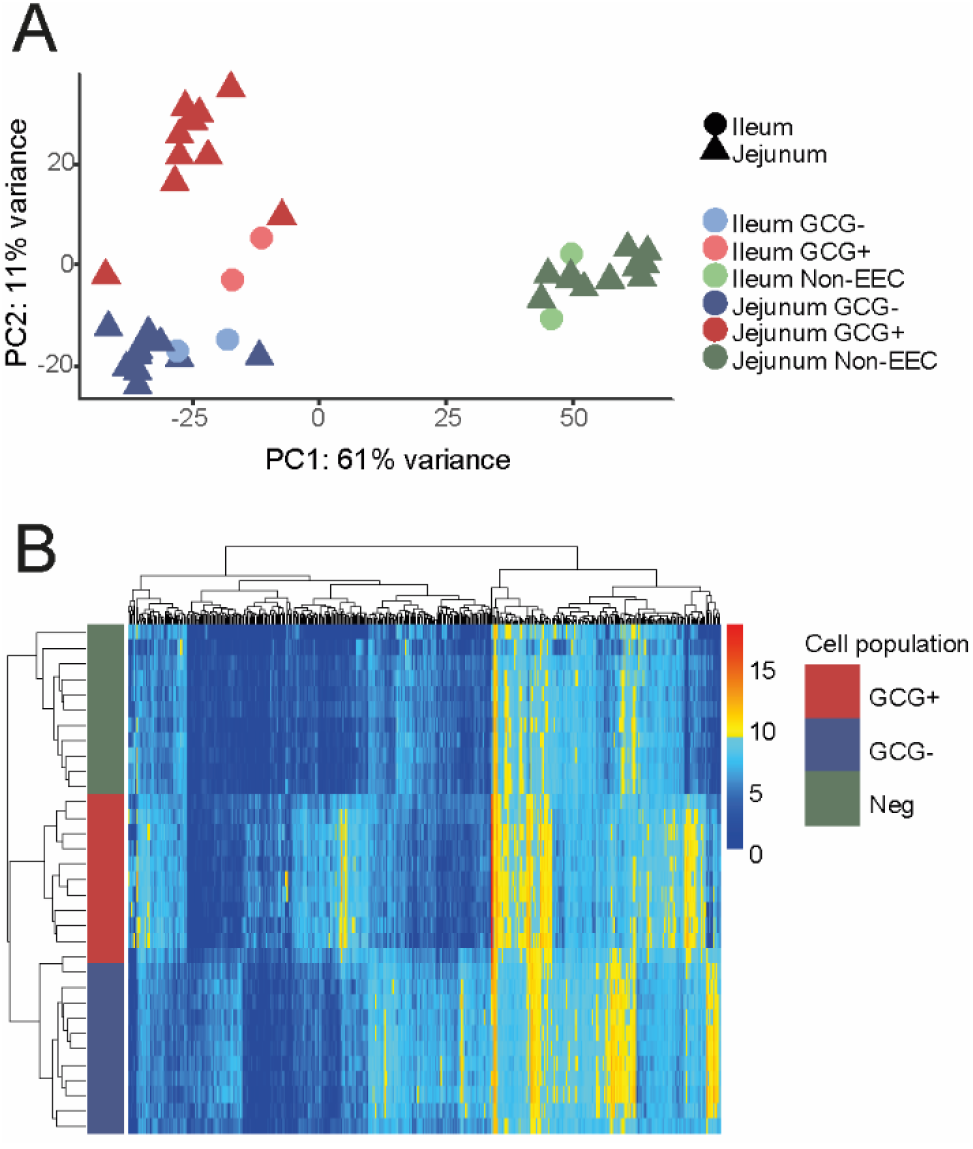
Transcriptomic distinction between cell populations from human small intestine. (A) Principal component analysis plot of all human samples (n=3 cell populations from each of 11 jejunal and 2 ileal tissue samples), differentiated by cell population and anatomical site. (B) Heatmap of 500 most differentially expressed genes in human jejunum samples (n=3 cell populations per each of 11 participants), y axis – cell population identified by coloured bar, x axis – genes.

Both human EEC populations expressed a wide range of hormonal transcripts (Figure 2A,C,E). As expected from the use of GLP-1 antibodies to purify GCG+ cells, the hormonal transcript showing the strongest differential expression in GCG+ vs GCG-samples was *GCG* itself (Figure 2C). Consistent with previous findings in mice(4, 12-14), we identified a range of additional hormonal transcripts in human GCG+ samples, including *GIP* (glucose-dependent insulinotropic polypeptide)*, CCK* (cholecystokinin), *NTS* (neurotensin), *PYY* and *SCT* (secretin) as well as *MLN* (motilin) – a hormone produced by human but not mouse(4, 12-15). Compared with GCG+ cells, GCG-cells had higher expression of *SCT*, *CCK*, *NTS*, *MLN*, *GHRL* (ghrelin) and *SST* (somatostatin), together with *TPH1*, the enzyme responsible for serotonin biosynthesis in enterochromaffin cells. EECs also expressed the putative gut hormones *UCN3* (urocortin 3), *PCSK1N* (ProSAAS) and *NPW* (neuropeptide W), as well as lower levels of RNAs encoding several peptides not classically described as gut hormones such as *VGF*, *GHRH* and *ADM*.

**Figure 2.**
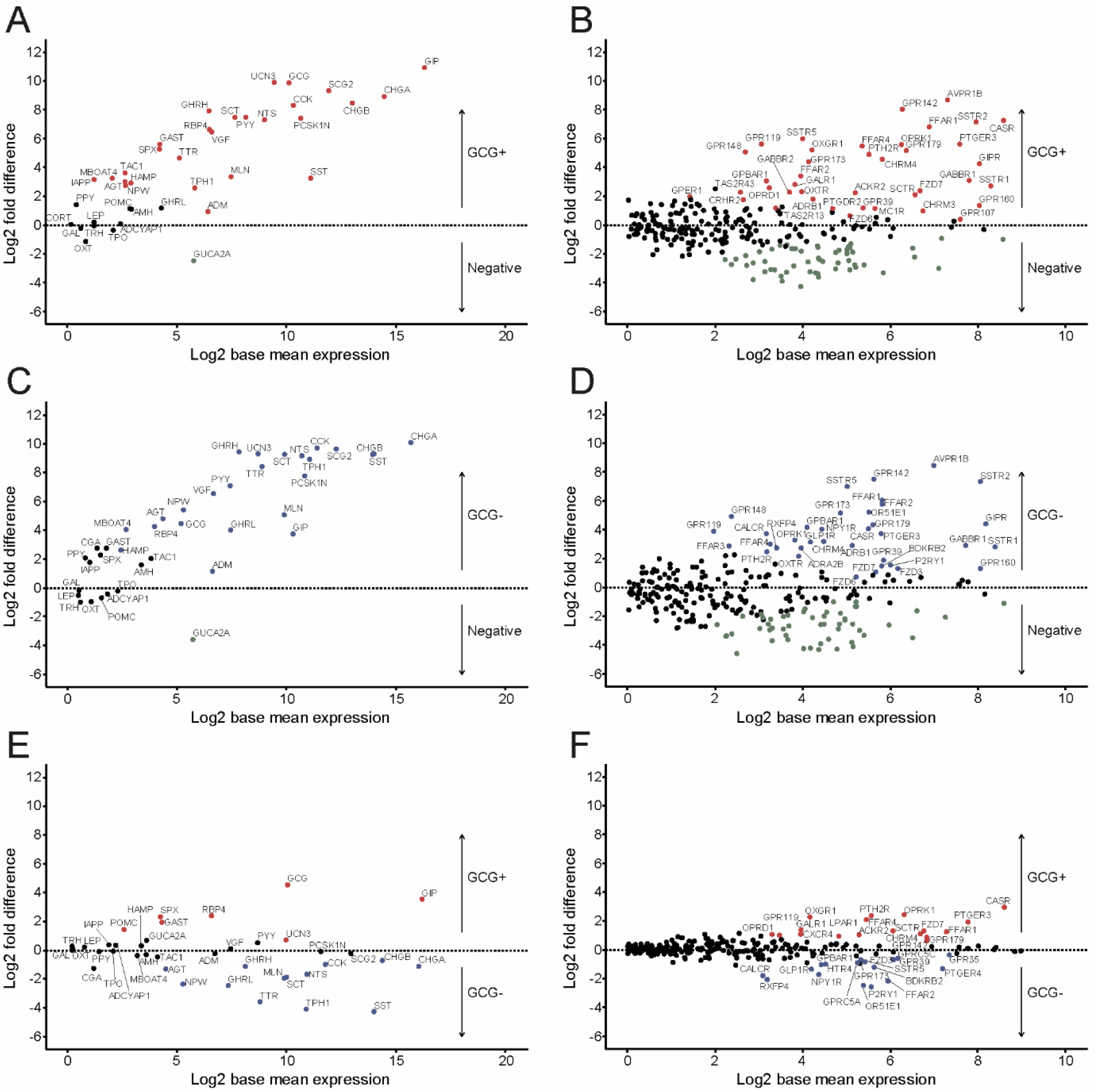
Transcripts enriched in human EECs. Enrichment vs expression plots for human jejunum EEC populations. Enrichment is presented as the log2 fold difference between the cell populations indicated, and expression is presented as the log2 base mean normalised expression extracted from the DESEQ2 model. (A,B) GCG+ vs negative, (C,D) GCG-vs negative, (E,F) GCG+ vs GCG-. Hormones and granins are shown in A,C,E; receptors and ion channels are shown in B,D,F. Red – enriched (adjusted P<=.1 in DESEQ2 model) in GCG+; Blue – enriched in GCG-; Green – enriched in negative cells.

Transcripts of ^~^50 GPCRs were either enriched (*P*<0.1) over the negative population, or expressed at > 100 CPM in one or both EEC populations (Figure 2B,D,F). Multiple GPCRs previously implicated in post-prandial gut hormone secretion in mice were highly expressed in human EECs compared with negative cells, including the fat-sensing receptors *FFAR1*, *FFAR2*, *FFAR3, FFAR4* and *GPR119,* the amino acid sensing receptors *CASR* and *GPR142,* the butyrate and isovalerate sensing *OR51E1*(16), and the bile acid receptor, *GPBAR1*, as well as the zinc receptor *GPR39,* and the alpha-ketoglutarate receptor *OXGR1.* Human EECs differentially expressed a number of hormonal receptors compared with negative cells, including *SSTR1*, *SSTR2, SSTR5*, *GIPR*, *AVPR1B, OXTR, CRHR2*, *RXFP4*, *GLP1R*, *NPY1R, CALCR, GALR1* and *SCTR*. Two opioid receptors, *OPRK1* and *OPRD1,* not previously described as active in the gastrointestinal tract were identified in EECs, alongside the oestrogen receptor *GPER1* and prostaglandin receptors *PTGDR2* and *PTGER3*. At least four orphan GPCRs were differentially expressed in human EECs: *GPR148*, *GPR160*, *GPR173* and *GPR179*, hinting to as yet undescribed pathways that may control gut hormone secretion.

Transcripts for a range of ion channel subunits were enriched in human EECs (Supplementary Figure 2), consistent with previous reports that murine L-cells and enterochromaffin cells are electrically active. Human GCG+ and GCG-EECs were particularly enriched for the voltage-gated calcium channel subunits *CACNA1A* (P/Q-type), *CACNA1C* and *CACNA1D* (L-type), and *CACNA1H* (T-type)(17), and the voltage-gated sodium channel subunits *SCN3A* and *SCN8A*. *TRPA1*, previously described to stimulate GLP-1 and 5-HT secretion in mouse, was enriched in both EEC populations(18, 19), as was the amiloride/acid sensitive ion channel *ASIC5*, and the K_ATP_ channel subunits *KCNJ11* and *ABCC8*(8, 20).

Transcription factor profiling of human GCG+ and GCG-cells is also shown in Supplementary Figure 2.

### Comparison of human jejunum vs ileum EEC transcriptomes

PCA of the 3 cell populations from each of the 2 matched human jejunum and ileum samples separated EECs from negative cells on the first component and the anatomical location of GCG+ and GCG-cells on the second component (Figure 3A). This distinction was driven by a low number of differentially expressed genes, 279 in GCG+ cells and 120 in GCG- cells, when analysed pairwise using a DESEQ2 model. Notable differentially expressed transcripts that were higher in jejunal than ileal EECs (GCG+ or GCG-) included *GIP*, *CCK*, *SST*, *MLN*, *SCT*, *GHRH*, *ASIC5*, and *TRPA*1; whereas transcripts higher in ileal than jejunal EECs included *GCG*, *NTS* and *TAC1* (Figure 3,B,C).

**Figure 3.**
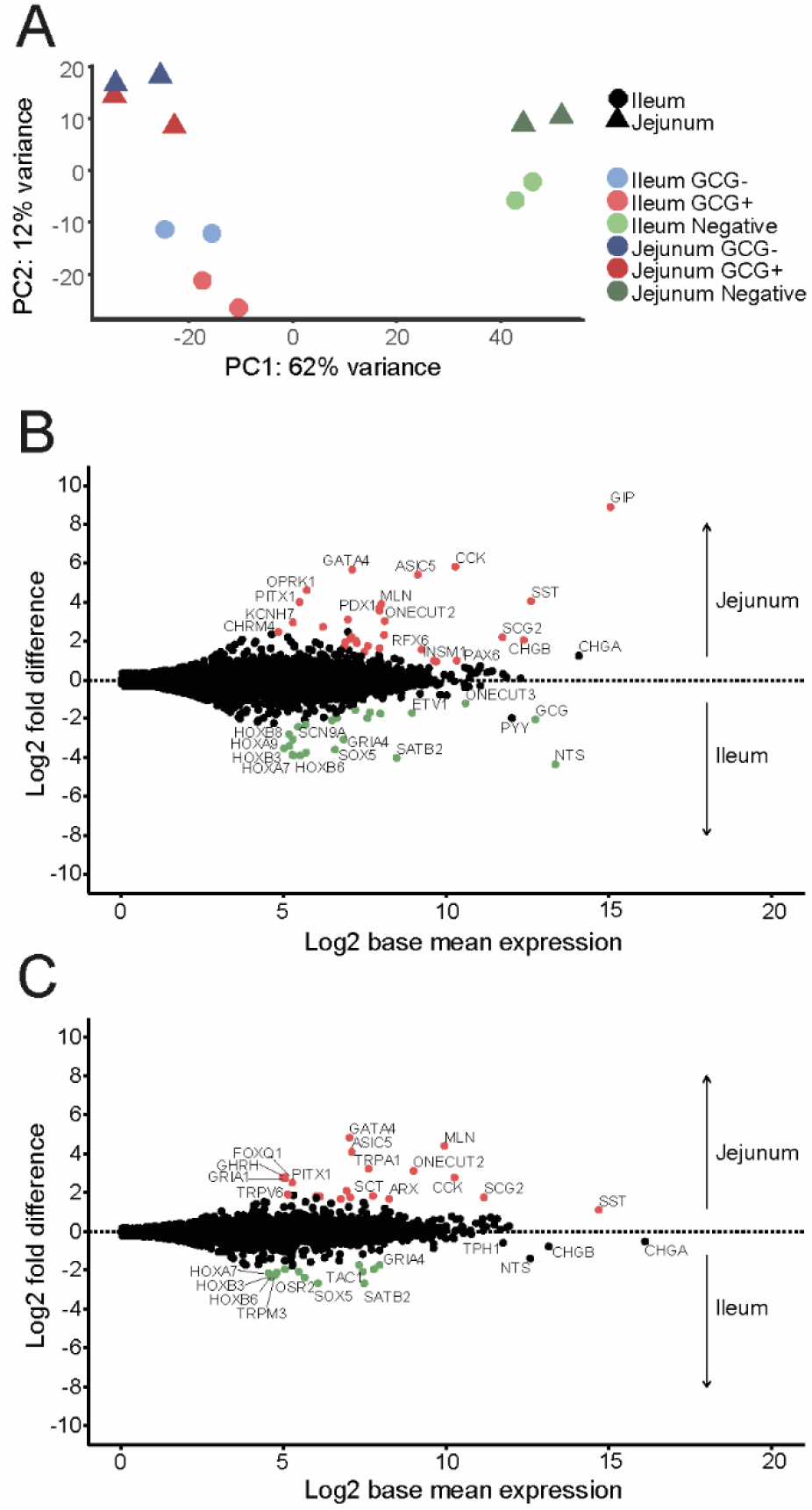
Comparison between EECs from human jejunum and ileum. (A) Principal component analysis plot of matched human jejunum vs ileum samples, labelled by cell population and anatomical location (n=3 cell populations from both anatomical regions from each of 2 participants). (B,C) Enrichment vs expression plots for GCG+ (B), and GCG- (C), cell populations. Enrichment is presented as the log2 fold difference between the cell populations indicated, and expression is presented as the log2 base mean normalised expression extracted from the DESEQ2 model. Red – enriched (adjusted P<=.1 in DESEQ2 model) in jejunum; Green – enriched in ileum.

### Comparison of human vs mouse EEC transcriptomes

We compared human jejunal GCG+ cells with murine upper small intestinal GLU-Venus cells, representing the GLP-1 secreting L-cell populations from each species (Figure 4). Only genes with 1:1 homology annotated in Ensembl were included, and genes annotated as ribosomal, mitochondrial and small-nuclear were excluded, giving a total of 15,507 genes for comparison. Log-log plots of normalised gene expression indicated a strong correlation between the two species (R^2^=0.73; Figure 4). Of the 354 genes lying outside the 99% confidence interval of the linear model of human versus mouse, notable genes more highly expressed in human than mouse GCG+ cells included *GIP*, *CHGA*, *ASIC5*, *GIPR*, *GPR142, SCTR, PTH2R, CHRNA5* and *OPRK1*; whereas genes more highly expressed in mouse GCG+ cells included *Gpr174*, *Gpr171, Ghr, Grpr, Ptger1, Cnr1, Insl5, Gpr22* and *Ghrl*.

**Figure 4.**
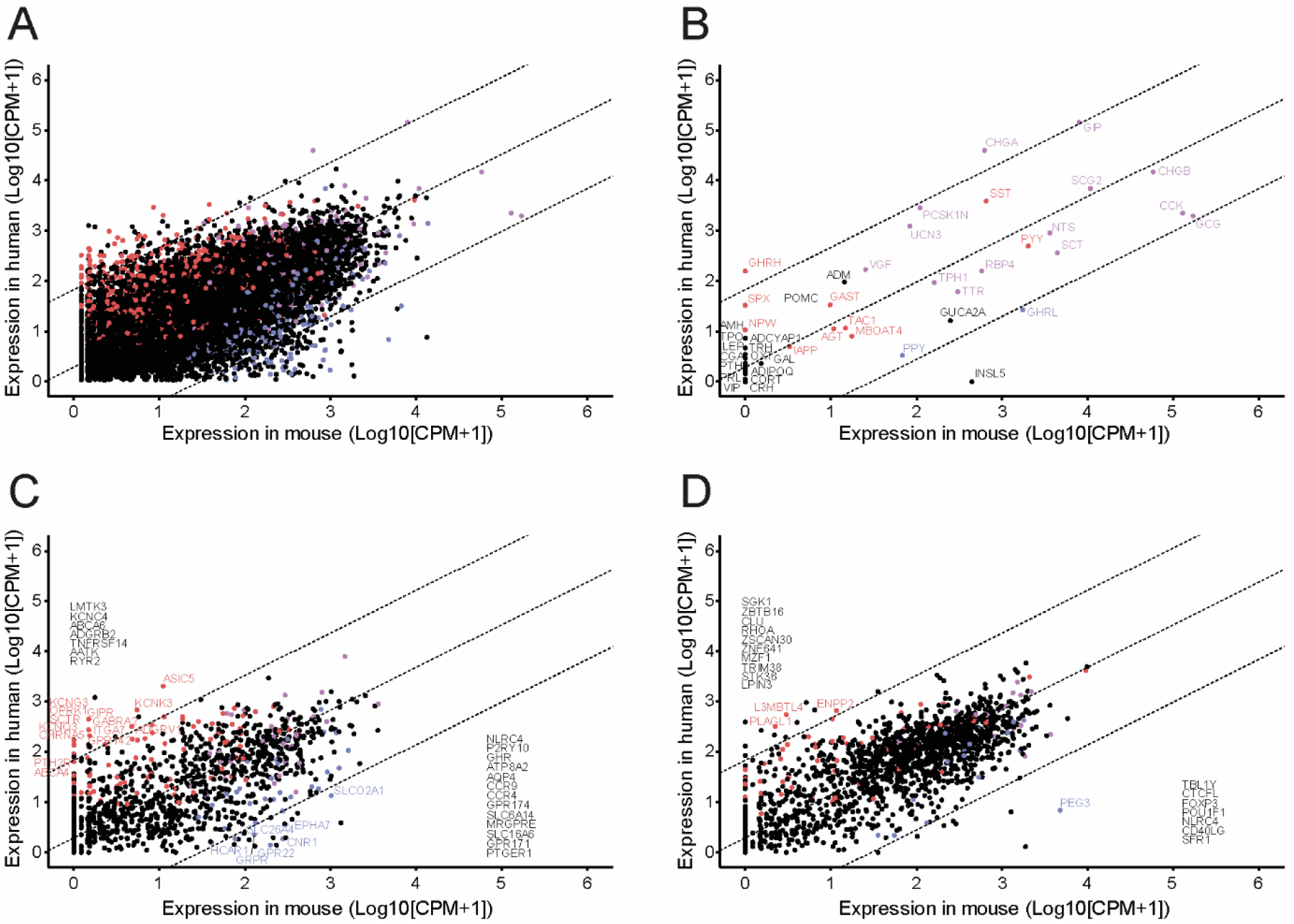
Comparison between human and mouse L-cells. Human versus mouse jejunal L cell gene expression (log10 normalised counts per million +1; n = 11 humans, n = 3 mice). (A) all genes with 1:1 homology between species, excluding mitochondrial, ribosomal and small nuclear transcripts (n=15,507). (B) Hormones. (C) GPCRs and ion channels. (D) Transcription factors. Dashed lines are linear regression and 99% confidence interval. Each dot represents normalised CPM+1 for one gene. Red – enriched (>4x fold change) and differentially expressed (adjusted P<=.1) for human GCG+ vs Negative cell populations, but not murine GLU-Venus vs Negative cell populations in relevant DESEQ2 model. Blue – enriched and differentially expressed for murine GLU-Venus vs Negative cell populations, but not human GCG+ vs Negative cell populations. Purple – enriched and differentially expressed in both murine GLU-Venus and human GCG+ cells versus relevant negative cell populations. Black – not enriched and differentially expressed in either human GCG+ or murine GLU-Venus cells versus relevant negative cell populations. All genes are labelled in B, and genes outside the 99% CI are labelled in C and D, with those not differentially expressed or enriched in either population listed along the axis.

Human jejunal GCG-cells, representing the wider murine EEC population (although depleted of L-cells) were compared with murine NeuroD1-positive cells, revealing a strong correlation between these human and murine EEC populations (R^2^=0.74; Figure 5). Of the 380 genes lying outside the 99% confidence interval of the linear model for this comparison, notable genes more highly expressed in human than murine EECs included *GIPR*, *SCTR,, GHRH, OPRK1, PTH2R* and *TRPV6*, and in murine compared with human cells included *Ghrl*, *Tac1*, *Iapp*, *Gpr22, Gast, Ptger1, Grpr* and *Ghr*.

**Figure 5.**
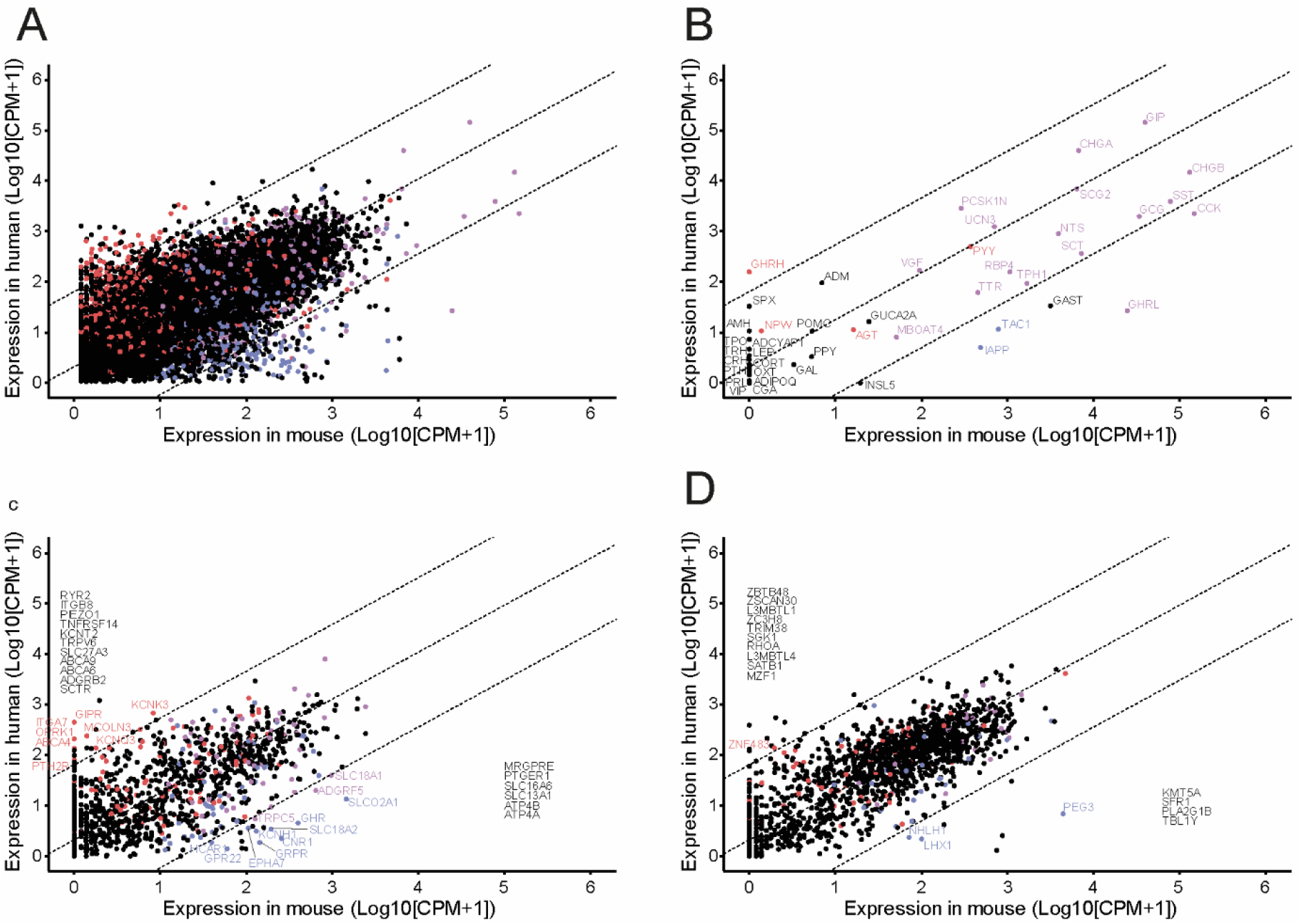
Comparison between human GCG- and mouse NeuroD1 cells. Human versus mouse jejunal EEC (GCG-) gene expression (log10 normalised counts per million +1; n = 11 humans, n = 3 mice). (A) all genes with 1:1 homology between species, excluding mitochondrial, ribosomal and small nuclear transcripts (n=15,507). (B) Hormones. (C) GPCRs and ion channels. (D) Transcription factors. Dashed lines are linear regression and 99% confidence interval. Each dot represents normalised CPM+1 for one gene. Red – enriched (>4x fold change) and differentially expressed (adjusted P<=.1) for human GCG- vs Negative cell populations, but not murine NeuroD1 vs Negative cell populations in relevant DESEQ2 model. Blue – enriched and differentially expressed for murine NeuroD1 vs Negative cell populations, but not human GCG- vs Negative cell populations. Purple – enriched and differentially expressed in both murine NeuroD1 and human GCG-cells versus relevant negative cell populations. Black – not enriched and differentially expressed in either human GCG- or murine NeuroD1 cells versus relevant negative cell populations. All genes are labelled in B, and genes outside the 99% CI are labelled in C and D, with those not differentially expressed or enriched in either population listed along the axis.

Species-comparative data for subgroups of genes in L-cells and the total EEC population, separated by their annotated roles as hormones, GPCRs, ion channels and transcription factors, are represented in Figures 5 and 6, in which the colour code additionally indicates whether the genes were more than four fold enriched and significantly likely to be differentially expressed based on the DESEQ2 model (adjusted P <= 0.1) in EECs compared with negative cells in one or both species.

**Figure 6.**
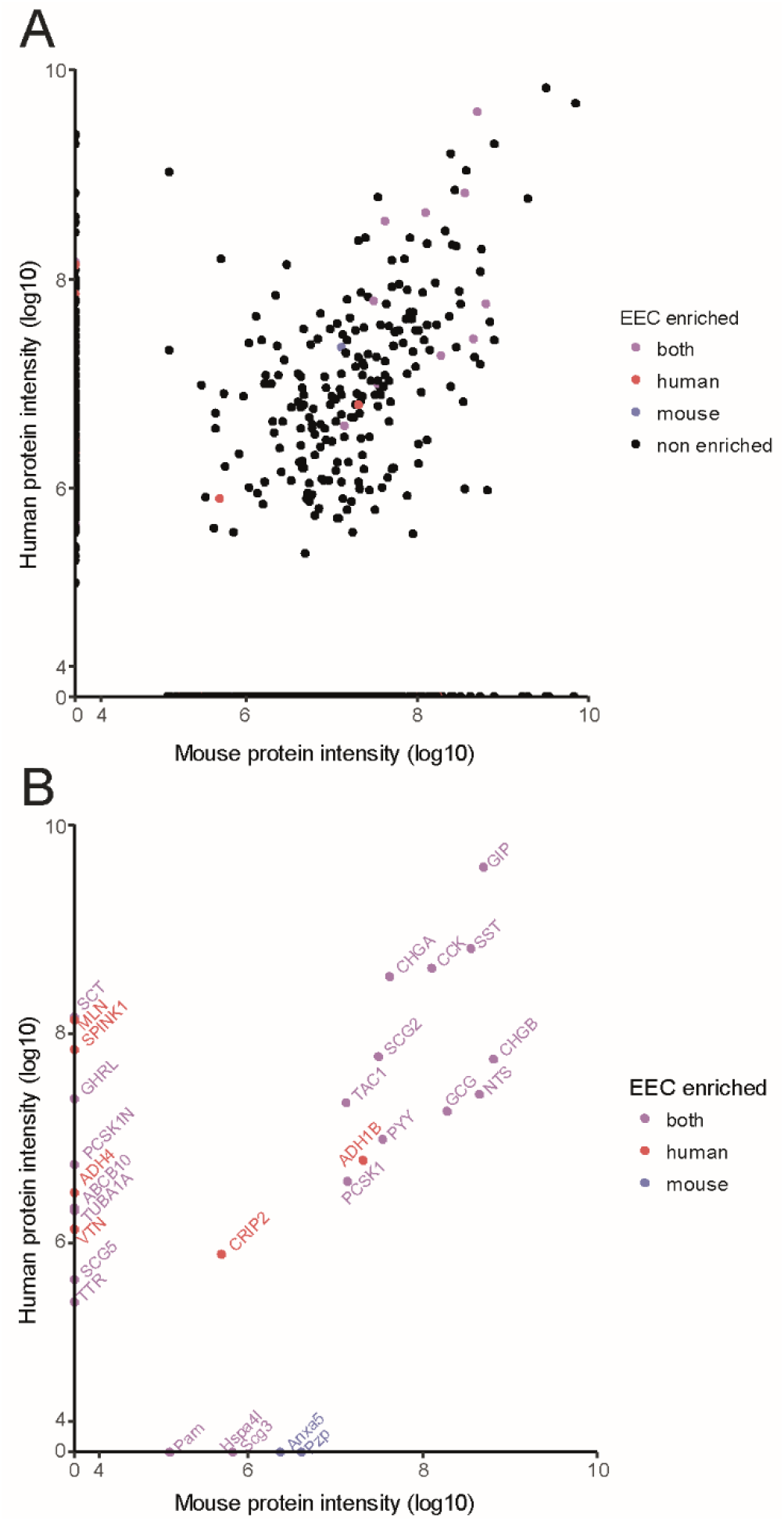
Comparison between human and mouse jejunum peptidome. Protein intensity as calculated by Peaks v8.0 for all proteins detected corresponding to genes with 1:1 homology between human and mouse for jejunum mucosal homogenates. (A) All proteins, (B) proteins from genes enriched in at least one of the species from the human and the mouse transcriptome datasets. Enrichment was defined as adjusted P<=.1, fold difference >4 and base mean expression >50 from DESEQ2 model. Colours indicate in which species the mRNA for the genes was enriched.

### Peptidomic analysis by LC-MS

Peptide extraction and LC-MS protocols for analysis of fresh intestinal mucosal samples were optimised for maximum peptide retrieval and identification (Supplementary Figure 3), and enabled reliable detection and sequencing of peptides up to 65 amino acids in length. In the first instance, tissue samples were analysed by nano LC-MS/MS and compared between human jejunum (n=4) and mouse mid-small intestine (n=4). Peptides were assigned to their parental proteins by Peaks software, and were found to include known EEC prohormones, granins and enteric neuronal signalling peptides as well as peptides derived from a variety of house-keeping proteins that likely reflected the occurrence of some tissue damage/degradation prior to homogenisation. Of the 463 and 705 different proteins matched in human and mouse respectively, 234 were common between the two species, and showed good correlation (R^2^ = 0.54, Figure 6A). To identify candidate EEC-derived peptides, we restricted the analysis to peptides originating from genes that in the transcriptomic analysis showed >4-fold higher expression in at least one EEC sample compared with the corresponding negative cell population (Figure 6B). The known gut hormone genes and members of the chromogranin family were mostly common to mouse and human, but motilin and ghrelin were found in human but not mouse jejunum, which is interesting, as the transcriptomic data identified *GHRL* to be preferentially expressed in mouse over human jejunum. A few peptides were derived from proteins not previously known to have signalling roles, but further studies will be needed to determine whether these exhibit bioactivity or simply reflect peptides released from EECs during tissue degradation. To search for novel candidate peptide hormones, we also examined the transcriptomic data for unannotated transcripts that had a base mean value >100 and were >16 fold more highly expressed in EECs than control cells. This analysis identified MIR7-3HG(21), C1orf127 and C6orf141, but we were unable to detect corresponding peptides in the LC-MS data.

We then performed a longitudinal LC-MS analysis of known bioactive peptides along the length of the mouse and human GI tract. Sequential samples were taken at 5cm intervals from the stomach to the rectum in mice (n=4 for each location), and human biopsies were analysed from the stomach (n=5), duodenum (n=9), jejunum (n=2), ileum (n=4), proximal colon (n=3), sigmoid colon (n=5) and rectum (n=3). Most EEC peptides were identifiable in their known bioactive forms but fragments of pro-CCK mostly lacked the C-terminally active 8 amino acids, perhaps reflecting an extraction artefact as CCK is known to be highly labile under different conditions and our method was not optimized for very small peptides such as CCK-8. We therefore used a robustly present peptide from the prohormone, CCK_21-44_, as a surrogate of CCK production. Peptides were depicted in separate heatmaps for mouse and human (Figure 7), divided into their origin from EEC prohormones, granins and non-EECs (likely reflecting enteric neural peptides). The observed longitudinal profiles broadly mirror historical immuno-staining patterns(22) but additionally provide details of the exact peptide sequences and their post-translational modifications that cannot be deduced from antibody staining. The human and mouse profiles were similar for most EEC peptides, with the notable exceptions that CCK, NTS and SCT extended into the colon only in mice, and that Gastrin and Ghrelin could be detected in human but not mouse proximal small intestine. Of the non-EEC peptides, NMU and galanin were found along the full length of mouse small and large intestine but in human NMU was largely restricted to the small intestine and galanin to the distal small intestine and colon/rectum.

**Figure 7.**
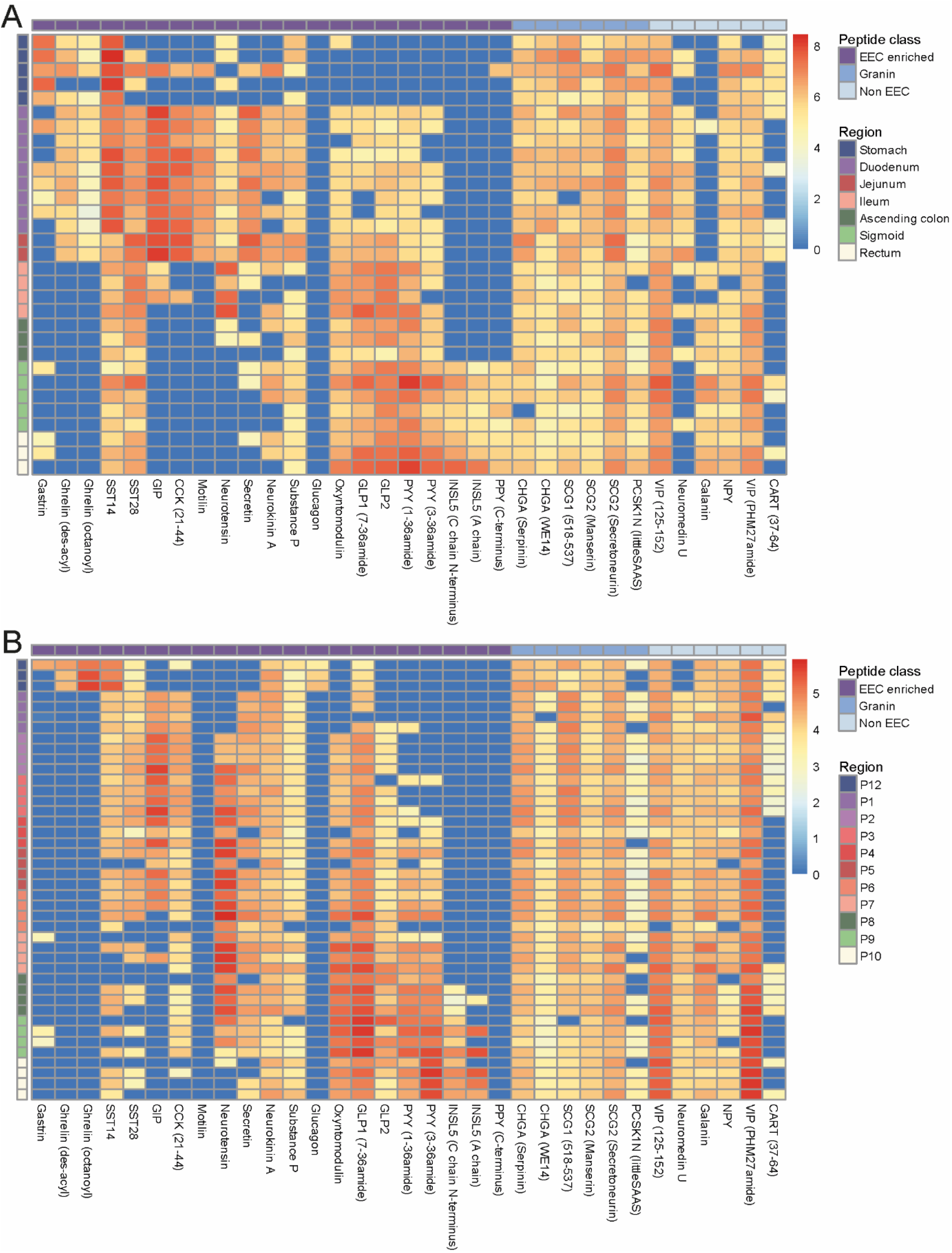
Longitudinal profiles of gut peptides along the human and mouse gut. Heatmap of gut peptide quantification normalised by tissue weight (log10 scale) along human (A) and mouse GI tract. Blue indicates not detected in the sample. Rows: Samples ordered from proximal to distal and colour coded by their region of origin. Mouse regions: P12: stomach lesser curvature, P1 to P7: small intestine from proximal to distal, sampling every 5 cm, P8 to P10: large intestine (proximal, mid and distal). Columns: peptide (using the human name if different between human and mouse), classified by origin (purple: classical EEC hormone peptides; medium blue: peptides from granins; light blue: enteric neuron bioactive peptides).

## Discussion

By RNA sequencing of fixed, FACS-purified cells from the human gut, we report here the transcriptome of human L-cells and the wider EEC population, and a between-species comparison showing a strong similarity with matching cells from the mouse. LC-MS based peptidomic analysis revealed longitudinal gradients of a range of EEC peptides, including their sequences and post-translational modifications, along the length of the human and mouse GI tract. The amalgamated transcriptomic and peptidomic picture provides a unique insight into the sensory apparatus and hormonal secretory profile of human EECs, and the extent to which the mouse is a valid model for studying the human enteroendocrine system.

The study is limited by many of the same challenges that have consistently hindered RNA and peptidomic analyses. We relied on good quality internal epitopes to purify human EECs for transcriptomics, but this required cell fixation and permeabilisation prior to cell separation by flow cytometry. The available RNA extraction and sequencing kits were only just technically capable of dealing with PFA-modified RNA and although the resulting high throughput sequencing consequently had a lower than desired mapping rate the read depth was sufficient for detection of low abundance mRNAs such as those encoding receptors and ion channels. The transcriptomes of human L-cells and GCG-EECs mapped robustly onto the homologous RNAseq data from freshly purified matching murine cell populations. Genes without a 1:1 homology between species or that were not protein encoding were excluded from the comparative analysis, but can be examined in the raw deposited datasets. An alternative transcriptomic approach we could have used that would also not depend on the availability of genetically labelled models is single cell RNA sequencing of large numbers of dissociated cells, as recently applied to murine intestinal organoids(10). Although EECs accounted for only a small percentage of the total cell count, they were readily identifiable from their transcriptomic signatures(10). This approach could be extended to human organoids or fresh human tissue, but the potential influence of prolonged organoid culture conditions on the cell transcriptome still needs to be established and the outputs of most single cell RNA sequencing methods are currently limited by a read-depth that precludes reliable quantification of low-abundance transcripts.

Although we separately RNA sequenced jejunal samples from 11 humans with varying BMI, we were unable to detect any effect of BMI on the EEC transcriptome. Tissue samples, were, however, collected and sorted in both France and the UK where pre-operative dietary intakes were likely quite different. As the high BMI samples largely originated from the French site and we accounted for the donor location when constructing the models of enriched transcripts, the study was insufficiently powered to detect small differences attributable to obesity. We cannot therefore exclude the possibilities that altered EEC function might arise as a result of obesity or variation in dietary intake, or contribute to the causation of obesity in a subset of people.

GPCRs that are strongly and specifically expressed in EECs underlie some of the key sensory roles of these cell types and deserve consideration as candidate drug targets for enhancing gut hormone secretion as a therapy for type 2 diabetes and obesity. The GPCR repertoire of human EECs largely mirrored their well-studied murine counterparts(9, 23-25), including expression of receptors for amino acids (*CASR*, *GPR142*), triglyceride digestion products (*FFAR1, FFAR4, GPR119*) and bile acids (GPBAR1), as well as for hormones such as somatostatin (*SSTR1, SSTR2, SSTR5*), GIP (*GIPR*) and arginine vasopressin (*AVPR1B*). Interestingly, *GPR142* was highly expressed in human EECs, supporting current studies looking to exploit its ability to stimulate both insulin and incretin hormone secretion(24).

RNAseq analysis revealed expression in human and mouse EECs of genes encoding known gut peptides, as well as several putative hormones previously described in the gut or elsewhere. Of the hormone-encoding genes with a 1:1 homology between species, there was a good correlation between the expression profiles of mouse and human. Motilin was expressed in human but not murine EECs, consistent with previous literature(15). Our optimised peptide extraction protocol combined with nano-LC-MS analysis enabled identification of the exact peptide sequences biosynthesised in human and mouse intestinal mucosa, including post-translational modifications, for peptides from ^~^8-10 to a maximum length of 65 amino acids. From the proglucagon gene, for example, we detected multiple processed and pre-processed products, including GRPP, oxyntomodulin, GLP-1_7-36 amide_, GLP-1_7-37_, GLP-1_1-37_, IP_131-142_, IP-GLP2 and GLP-2. Intact (pancreatic-type) glucagon was detected in samples from the mouse stomach, but was undetectable in the remainder of the intestine and colon from both species, conflicting with recent suggestions that the small intestine can secrete intact glucagon(26), but consistent with our own recent findings that post-prandial glucagon concentrations were not altered following gastric bypass surgery in lean subjects despite dramatic increases in GLP-1(27). Whether the gut might adapt to produce glucagon in pathological conditions such as obesity or type 2 diabetes cannot be concluded from our data. In addition to the set of well-recognised gut hormones, LC-MS also identified some additional peptides encoded by EEC-enriched genes, including peptides derived from PCSK1N, chromogranins and secretogranins. Whether any of these have specific physiological roles or are simply inactive by-products of enzymatic processing of the contents of secretory vesicles, requires further evaluation.

Mapping of gut hormone production along the GI tract length has previously been performed using immuno-staining methods and extraction/immuno-assays for specific peptides(22, 28). Immuno-staining relies on the detection of all antigenic sequences binding to a primary antibody and is a reliable method for localising cells producing any given prohormone but cannot generally distinguish whether the prohormone was processed or unprocessed, or post-translationally modified. Nevertheless, immunostaining methods have resulted in longitudinal maps of EEC sub-types producing different prohormones(22). Our LC-MS method provides a surprisingly robust mirror of these immuno-staining maps, whilst assigning an exact peptide sequence to each identified peptide, clearly distinguishing e.g. oxyntomodulin from glucagon, and PYY_1-36_ from PYY_3-36_, despite these peptide pairs differing only by extensions of a few amino acids at the C and N-terminus, respectively. The method also benefits from being unbiased, providing MS-based identification of all peaks triggered from the liquid chromatography separation, rather than analysing only a sub-group of specified peptide sequences. Interestingly we identified acylated as well as non-acylated ghrelin from the human jejunum despite our previous finding that plasma acylated ghrelin levels were undetectable in humans after total gastrectomy(27). We were surprised to find high levels of PYY_3-36_ as well as PYY_1-36_ in tissue homogenates, suggesting that dipeptidyl-peptidases (DPP) are active within L-cells themselves, whereas GLP-1(7-36amide) was much more abundant than GLP-1(9-36amide), indicating they had not been subject to DPP cleavage. Why GLP-1 but not PYY seems protected from DPP-mediated processing in L-cells, despite both peptides being located in the same vesicular pool (manuscript under preparation), remains unclear.

In summary, we have developed techniques for RNA sequencing of purified human EECs, and for peptidomic analysis of mucosal homogenates using LC-MS. Comparison of the human and mouse EEC transcriptomes revealed strong global similarities between the two species. Variation at the level of a few key individual genes could, however, have profound implications for the use of mouse as a model species in drug development programmes targeting specific receptors. Our mouse/human comparative datasets provide tools for confirming the validity of using mouse as a model for investigating given signalling pathways and receptors or peptide processing in humans, and the human EEC GPCR-ome can be used independently as a potential source of drug targets in the human enteroendocrine system. Longitudinal peptide mapping of the GI tract by LC-MS is a key step towards understanding the metabolic benefits of gastric bypass surgery, as the surgical procedures result in a shift in the location of nutrient digestion and absorption to more distal regions of the small intestine, with consequent stimulation of the more distal EEC population and release of their distinct profiles of peptide hormones. Devising strategies to mimic gastric bypass surgery using injectable peptide mimetics or through stimulation of the endogenous EEC population is a key but highly topical challenge for industry and academia alike.

## Methods

### Ethics

This study was conducted in accordance with the principles of the Declaration of Helsinki and good clinical practice. Human ethical approvals were given by Cambridge Central and South Research Ethics Committees (ref: 09/H0308/24, 16/EE/0338, 15/EE/0152) and the Inserm ethics committee and Agence de la biomédecine (ref: PFS16-004). Animal work was regulated under the Animals (Scientific Procedures) Act 1986 Amendment Regulations 2012 and conducted following ethical review by the University of Cambridge Animal Welfare and Ethical Review Body.

#### Human tissue transcriptome

##### Sample collection

Samples of human jejunum discarded during surgery were collected during total gastrectomy for treatment or prophylaxis of gastric cancer, or Roux-en-Y gastric bypass for obesity. All were from the point of entero-enterostomy 50cm distal to the ligament of Treitz. Two matched samples of jejunum and terminal ileum were collected during organ procurement from transplant donors. Data were collected on age, gender and BMI, and participants stratified as lean vs obese (BMI>30kg/m^2^). Tissue samples from different regions of the human GI tract for LC-MS were obtained from Addenbrooke’s Human Research Tissue Bank and the Cambridge Biorepository for Translational Medicine.

Samples were immediately placed in cold Leibovitz’s L-15 media (Thermo Scientific, Waltham, MA, USA) and processed to the point of fixation or homogenised and stored at −70°C within 6 hours.

##### Tissue preparation for FACS

FACS and RNA extraction from fixed human cells followed a modified version of the MARIS protocol(29). Intestine was rinsed in cold phosphate buffered saline (PBS) and the muscular coat removed. Diced mucosa was digested twice in 0.1% w/v collagenase XI (Sigma-Aldrich, MO, USA) in Hanks’ Buffered Saline solution (HBSS) #9394 (Sigma-Aldrich) for 30 minutes each, shaking vigorously every 10 minutes. Supernatants were triturated, passed through a 50μm filter and centrifuged at 300g. Pellets were resuspended in PBS and fixed in 4% w/v paraformaldehyde (PFA) at 4°C for 20 minutes. PFA-fixed cells were washed twice in nuclease free 1% w/v bovine serum albumin (BSA) in PBS, and if a FACS facility was not immediately available, were suspended in 1% w/v BSA and 4% v/v RNAsin plus RNAse inhibitor (Promega, WI, USA) in PBS at 4°C overnight.

Cells were permeabilised with either a single 30 minute incubation with 0.1% v/v Triton x100 (Sigma-Aldrich) in 1% w/v BSA in PBS prior to antibody staining, or by the addition of 0.1% w/v Saponin (Sigma-Aldrich) to solutions in all steps from this point until after the first wash post-secondary antibody staining, with identical results.

Primary antibody staining was for one hour in 4% v/v RNAsin, 1% w/v BSA, 1% v/v goat anti-GLP-1 (Santa Cruz, Dallas, TX, USA; sc7782), 2% v/v rabbit anti-CHGA (Abcam, Cambridge, UK; Ab15160), 0.25% v/v rabbit anti-SCG2 (Abcam, Ab12241) in PBS at 4°C. Cells were then washed twice in 1% w/v BSA, 1% v/v RNAsin, and secondary antibody staining was for 30 minutes in 4% v/v RNAsin, 1% w/v BSA, 0.2% v/v donkey anti-goat Alexa 555, 0.2% v/v donkey anti-rabbit Alex 647 in PBS at 4°C. Cells were washed twice then suspended in 4% v/v RNAsin, 1% w/v BSA in PBS on ice for FACS.

##### FACS

Cell populations were sorted on a BD FACS ARIA III in the Cambridge NIHR BRC cell phenotyping hub or at Institut Cochin, Paris. Single cells positive for Alexa 647 but not Alexa 555 (i.e. CHGA/SCG2 +ve / GLP-1 –ve) were classified as GCG- enteroendocrine cells. Single cells positive for both Alexa 647 and Alexa 555 were classified as GCG+ enteroendocrine cells. At least 5000 cells were collected for each positive population. Twenty thousand double negative cells were collected as the negative (i.e. non-enteroendocrine) cell population. Cells were sorted into 2% v/v RNAsin in PBS at 4°C.

##### RNA extraction

RNA was extracted using the Ambion Recoverall Total nucleic acid isolation kit for FFPE (Ambion, CA, USA) with modifications to the protocol as below. The FACS sorted cell suspension was centrifuged at 3000g for 5 minutes at 4°C and the pellet resuspended in 200μl digestion buffer with 4μl protease and incubated at 50°C for 3 hours. The solution was then stored at −70°C for at least 12 hours prior to further extraction. After thawing, RNA was extracted using the manufacturer’s protocol with the exception of performing 2x 60μl elutions from the filter column in the final step.

The RNA solution was concentrated using a RNEasy Minelute cleanup kit (Qiagen, Hilden, Germany). RNA aliquots were diluted to 200μl with nuclease free water. The standard manufacturer’s protocol was followed with the exception that 700μl, not 500μl, of 100% ethanol was added to the solution in step two, to generate optimum binding conditions for the PFA fragmented RNA. RNA concentration and quality was analysed using an Agilent 2100 Bioanalyser (Agilent, CA, USA).

##### Sequencing

cDNA libraries were created using the Clontech SMARTer Stranded Total RNA-Seq Kit – Pico Input Mammalian v1 (Takara Bio, USA). RNA input quantity was 5ng and the non-fragmentation protocol was used. The standard manufacturer’s protocol was followed with the exception that 175μl of AMPure beads were used for the final bead purification to ensure recovery of the small fragments of RNA arising from PFA fixation. Sixteen PCR cycles were used for amplification.

50 base single-end sequencing was performed using an Illumina HiSEQ 4000 at the CRUK Cambridge Institute Genomics Core.

#### Mouse transcriptome

##### Sample collection and preparation for FACS

Female NeuroD1-Cre/EYFP (mixed background: 3-10 generations back-crossed with C57BL6) and GLU-Venus mice (C57BL6)(7, 8)aged 8-10 weeks were killed by cervical dislocation (n=3 each). Diced mucosa from the proximal 10cm of small intestine was digested twice in 0.1% w/v collagenase XI in HBSS at 37°C for 30 minutes each. Cells were pelleted by centrifugation at 100g for 1 minute, triturated and passed through a 50μm filter. Cells were stained with DAPI (1μg/ml) for 5 minutes at room temperature, washed twice and sorted in HBSS on a FACSJazz sorter at the Cambridge NIHR BRC cell phenotyping hub.

##### FACS and RNA extraction

All positive cells, and 20,000 negative cells were collected separately into aliquots of 500μl of buffer RLT+ (Qiagen), with 143mM β-mercaptoethanol. RNA was extracted using a RNeasy Micro plus kit (Qiagen) and quantified using an Agilent 2100 Bioanalyser

##### Sequencing

2ng of each RNA was used for cDNA amplification by SPIA amplification using the Ovation RNAseq system V2 kit (Nugen, CA, USA). 1μg of cDNA was then fragmented to ^~^200bp by sonication (Diagenode, Liege, Belgium) and adaptors for the indexing were added using the Ovation Rapid DR multiplex 1-96 kit. Samples were pooled and concentrated together with a MinElute column (Qiagen) to reach a concentration of 10nM. Single end 50 base sequencing was performed at the CRUK Cambridge Institute Genomics Core with an Illumina Hiseq4000.

#### RNAseq pipeline

Quality control and trimming of adaptors was performed using FastQC(30). Sequenced transcripts were mapped to the human (GRCh37) and mouse (GRCm38) genomes using TopHat 2.1.0 and raw counts generated using Cufflinks 2.2.1(31-33). Differential gene expression analysis was performed in RStudio using DESEQ2(11). Gene annotation was pulled from the Ensembl dataset held in BioMart(31). Receptor and ion channel lists were generated from the IUPHAR “targets and families” list(34). Graphical output used ggplot2 and pheatmap in RStudio(35).

#### Comparative transcriptomics

Mouse and human data sets were compared using only the 15,507 genes present in both datasets, not annotated as ribosomal, mitochondrial or small-nuclear and described with one-to-one homology according to the Ensembl mouse-human homology dataset(31). Normalised CPM (counts per million) were generated from the respective DESEQ2 models and compared for the human GCG+ population versus the murine GLU-Venus population and the human GCG-population versus the murine NeuroD1 population. Linear models were generated of the log10 CPM of the human vs murine datasets by a total least squares strategy, 99% confidence intervals calculated and the outliers hand searched for relevant genes.

Examining just the human samples, DESEQ2 models were generated for the following sets of jejunum samples: GCG+ versus negative; GCG-versus negative; GCG+ versus GCG-; GCG+ lean versus obese; GCG-lean versus obese. Participant paired DESEQ2 analyses were also performed comparing GCG+ and GCG-populations from the jejunum and ileum of the two transplant donor participants, for whom there were matched jejunum and ileum samples. An adjusted P value of <=.1 was defined as the cut off for statistical significance when examining for differential gene expression between populations.

#### Tissue homogenates

Small pieces (^~^2×2mm, 15-45 mg) of mucosa from different regions of the human GI tract or from every 5cm of the GI tract from C57BL6 mice were homogenised in 250μl of 6M guanidine hydrochloride (Sigma-Aldrich) with Lyzing MatrixD (MPbio), in a FastPrep-24 for 4×40s at 6ms^-1^. Samples were stored at −70C before further processing. Proteins were precipitated by adding 80% acetonitrile in water then centrifuged at 12000g at 4°C for 5 minutes. The aqueous phase containing the peptides was collected, dried on a centrifugal vacuum concentrator and stored at −70°C before analysis

#### Mass spectrometry

Samples were extracted using a Waters HLB μElution solid-phase extraction (SPE) plate (Waters, MA, USA) after being resuspended in 500μL 0.1% v/v formic acid in water as described previously and analysed after reduction/alkylation(36). Human homogenates and mouse homogenates for interspecies comparison were analysed using nano-flow based separation and electrospray approaches on a Thermo Fisher Ultimate 3000 nano LC system coupled to a Q Exactive Plus Orbitrap mass spectrometer (ThermoScientific). High-flow separation for the longitudinal mouse analysis was as previously described(36). Downstream analysis was performed using Peaks 8.0 software (Waterloo, ON, Canada) against the human and the mouse Swissprot databases (downloaded 26^th^ October 2017)(37), with a fixed cysteine carbamidomethylation and variable methionine oxidation, N-terminal acetylation and pyro-glutamate and C-terminal amidation modifications. Manual searches were performed for other modifications. Peptides of interest were quantified by measuring peak areas for selected m/z ranges and retention times corresponding to the peptide sequences and normalised by tissue weight.

## Contributions

GR, PR, PL, RS, RH, FR and FG designed the study. Tissue handling and FACS were performed by GR, PR, PL and LG. cDNA library production and bioinformatics analysis were performed by GR, PR, PL, MM and BL with support from GY and DC. Mass spectrometry was by PL, RK, EM, SG and GR. NeuroD1 mice were generated by AL and HJL. The manuscript was written by GR, FR and FG. All authors have had editorial oversight of the final manuscript version.

## Acknowledgements

We would like to thank the surgical team in the Cambridge Oesophago-gastric centre, Sébastien Czernichow, Cyrille Ramond, The Cambridge Biorepository for Translational Medicine and the NIHR Cambridge Human Research Tissue Bank for assistance with collecting donor tissue. FACS was supported by the Institut Cochin in Paris and the Cambridge NIHR BRC Cell Phenotyping Hub. Metabolic Research Laboratories support was provided by the following core facilities: Disease Model Core, Genomics and Transcriptomics Core, Histology Core, Imaging Core and NIHR BRC Core Biochemical Assay Laboratory (supported by the MRC [MRC_MC_UU_12012/5] and Wellcome Trust [100574/Z/12/Z]). RNA-sequencing was undertaken at the CRUK Cambridge Institute Genomics Core.

GR received an Addenbrooke’s Charitable Trust / Evelyn Trust Cambridge Clinical Research Fellowship [16-69], an EFSD project grant and a Royal College of Surgeons Research Fellowship. SG is supported by a grant from the BBSRC. Research in the laboratory of Fiona Gribble and Frank Reimann is supported by the MRC [MRC_MC_UU_12012/3], Wellcome Trust [106262/Z/14/Z, 106263/Z/14/Z] and research grants from Medimmune. The MS instrument was funded by the MRC “Enhancing UK clinical research” grant (MR/M009041/1). PR was supported by a postdoctoral grant from Agence Nationale de la Recherche (Laboratoire d’Excellence Revive, Investissement d’Avenir; ANR-10-LABX-73). The RS lab is supported by ANR-10-LABX-73 and the Bettencourt-Schueller foundation. This work was partially supported by the following NIH grants to AL: R01 DK100223-02, R01 DK110614-01.

## Supplementary material

**Supplementary table 1.**
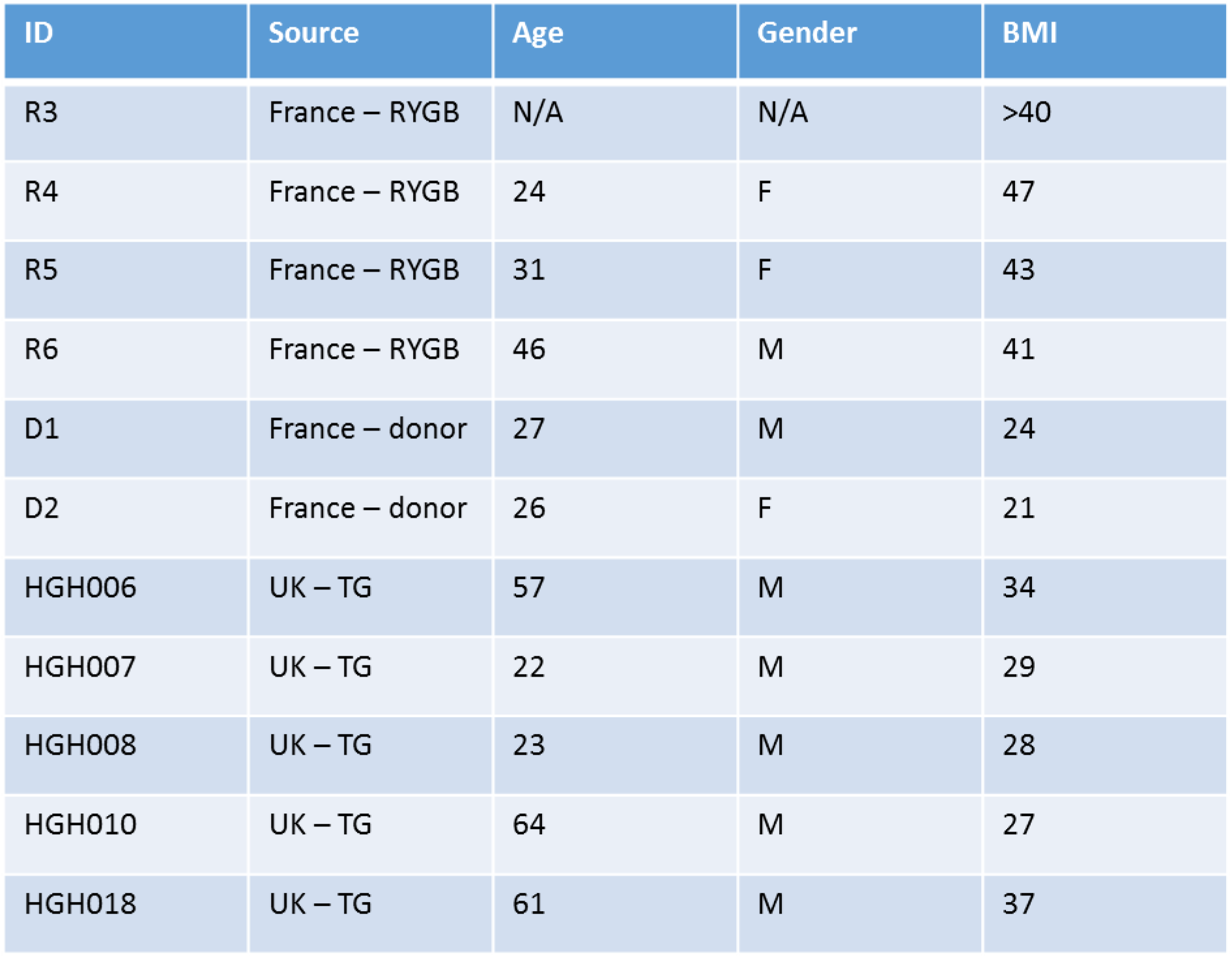
Demographics of human participants from transcripomic study. RYGB = Roux-en-Y Gastric Bypass; donor = organ transplant donor; TG = Total gastrectomy for prevention / treatment of gastric cancer

| ID | Source | Age | Gender | BMI |
| --- | --- | --- | --- | --- |
| R3 | France – RYGB | N/A | N/A | >40 |
| R4 | France – RYGB | 24 | F | 47 |
| R5 | France – RYGB | 31 | F | 43 |
| R6 | France – RYGB | 46 | M | 41 |
| D1 | France – donor | 27 | M | 24 |
| D2 | France – donor | 26 | F | 21 |
| HGH006 | UK – TG | 57 | M | 34 |
| HGH007 | UK – TG | 22 | M | 29 |
| HGH008 | UK – TG | 23 | M | 28 |
| HGH010 | UK – TG | 64 | M | 27 |
| HGH018 | UK – TG | 61 | M | 37 |

**Supplementary figure 1.**
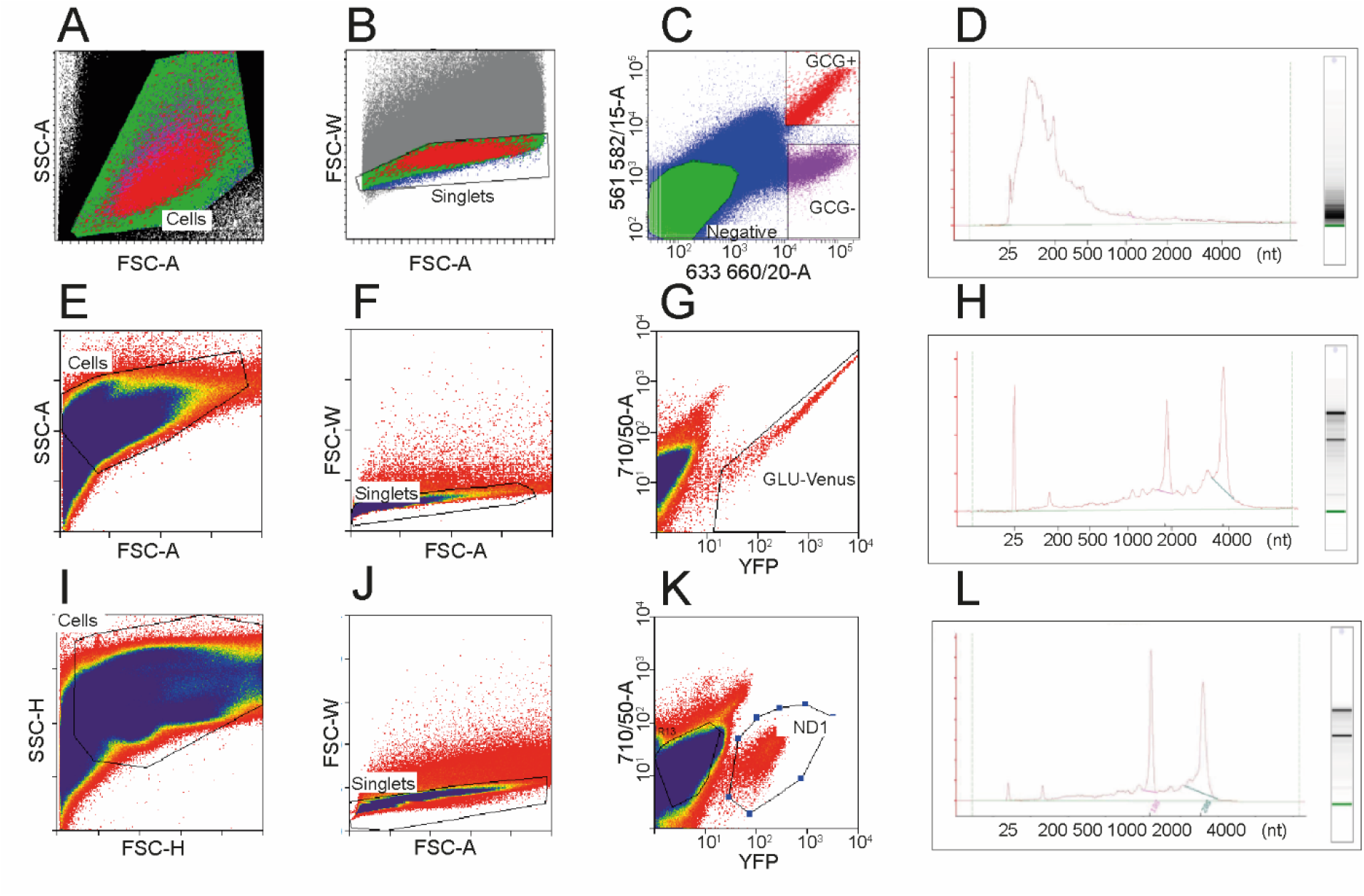
(A,B,C) FACS sorting of human GCG+, GCG- and negative cells. Cells were sorted by side scatter (SSC), forward scatter (FSC-A) and pulse width (FSC-W) to collect single cells. CHGA/SCG2 staining was detected by 633nm excitation, 660/20nm emission; GLP-1 staining was detected by 561nm excitation, 582/15nm emission. (E,F,G) FACS sort of murine GLU-Venus cells using gating as labelled. (I,J,K) FACS sort of murine NeuroD1-Cre/ROSA26-YFP cells using gating as labelled. (D,H,L) Agilent Bioanalyzer 2100 trace of RNA quality from human GCG+ (D) and murine GLU-Venus (H) and NeuroD1 (L) cell populations.

**Supplementary figure 2.**
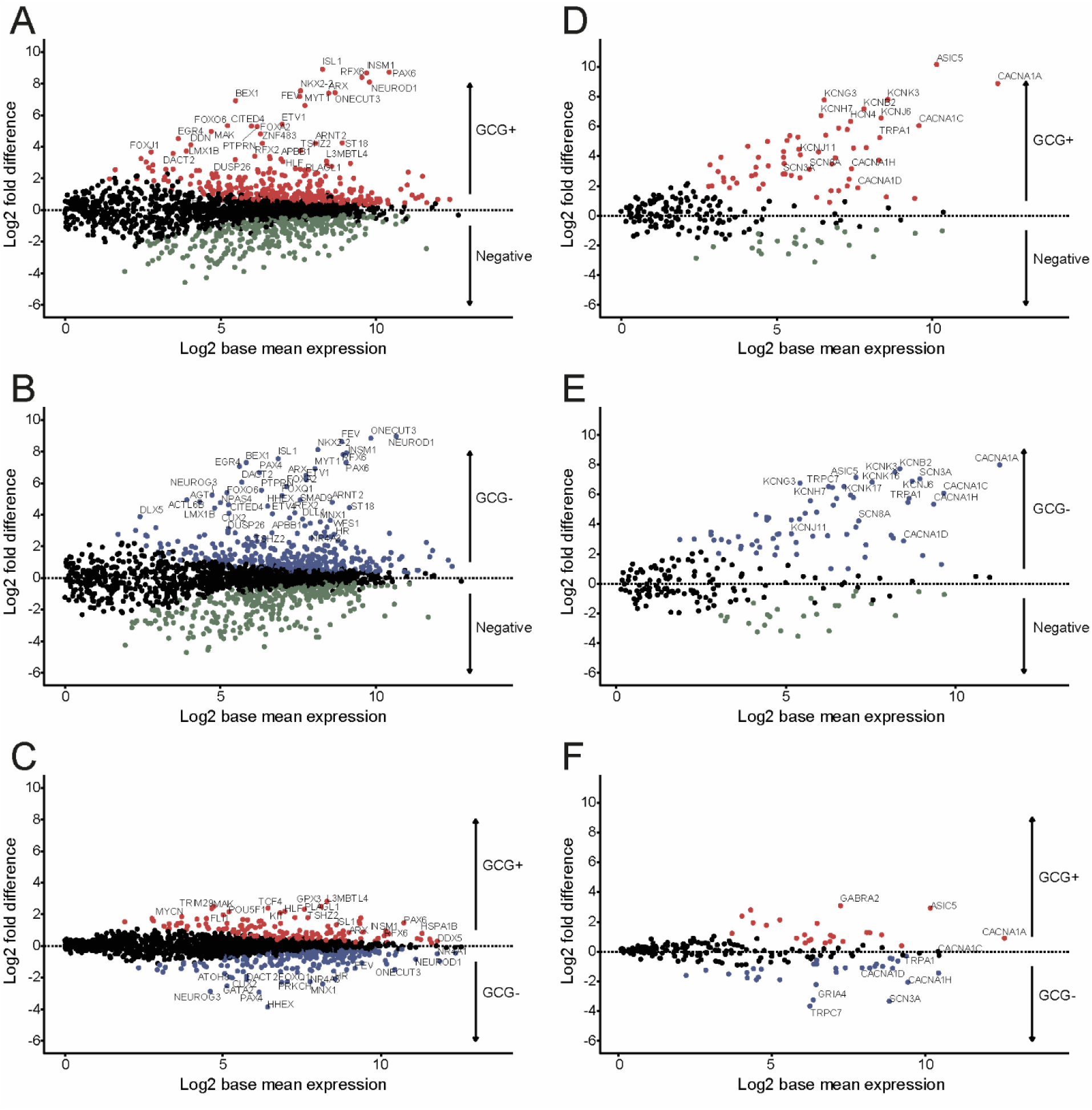
Differntial vs mean expression plots for human jejunum EEC populations. Differential expression is presented is presented as the log(2) fold difference between the cell populations indicated, and expression is presented as the log(2) base mean normalised expression extracted from the DESEQ2 model. (A,D) GCG+ vs negative (C,F), GCG+ vs GCG-, Transcription factors are shown in A,B,C; ion channels are shown in D,E,F. Red – enriched (i.e. adjusted p<0.1) in GCG+; Blue – enriched in GCG-; Green – enriched in negative cells. Key genes are labelled

### Supplementary figure 3 – Peptidomics optimisation studies

Peptidomics is a relatively new and under-developed field within proteomics, despite the varied and significant roles of peptides as signalling molecules in many organisms. Peptides, due to their small size in comparison to proteins, have specific properties that can be used to extract and analyse them intact by mass-spectrometry, giving information about their exact processing and post-translational modifications. Several methods to extract peptides from different tissues or fluids have been published, but none was developed to study intestinal tissues and the large variety of hormone peptides they produce.

Peptide extraction and purification for LC-MS/MS studies initially requires the complete disruption of the tissue of interest while inhibiting all protease activity. This is to avoid contamination with degradation products of larger proteins as well as degradation of the peptides of interest. The low-molecular weight proteins and peptides must then be separated from the rest of the proteins and other molecules. We compared several methods adapted from the literature to develop a robust and reliable method to quantify peptide levels from intestinal tissues. Methods tested for tissue lysis and protein extraction included tissue homogenization in 0.25% acetic acid, 8M urea, 90% methanol with 1% acetic acid, guanidine hydrochloride or 80% acetonitrile, or using Trizol to extract both proteins and peptides from the same sample^1–4^ Peptides were separated from proteins using either 10kDa molecular weight cut-off filters or 80% acetonitrile. Samples were finally purified using SPE columns, dried and reduced alkylated before being analysed by mass-spectrometry.

The different methods allowed the detection of many unique peptides by LC-MS/MS with an important variability between the different methods, from 1778 unique peptides for the extraction using 8M urea to 25 with 90% methanol/1% acetic acid. Many corresponded to fragments of longer peptides or proteins and could represent degradation products, coming either from the protein turn-over of the tissue or degradation happening during sample preparation. However, most of the peptides were unique to the extraction method (Supp Fig 2a), due to their different chemical properties and also due to different degradation levels. Focusing on the analysis of the classical gut peptide hormones significantly reduced the number of hits, and clearly showed that most of the expected intact peptides were detected using the guanidine HCl extraction method (Supp Fig2b). In addition to improving the number of detected hormone peptides, the guanidine HCl extraction method showed the highest levels of these peptides when quantified (Supp Fig 2c). Urea and Trizol extraction were the second best methods for peptide levels and variety of peptide hormones extracted, and tissue homogenization using Trizol can be a reliable technique to extract both RNA and peptides from a same sample. We also compared peptide extraction from the proteins using either protein precipitation with 80% ACN or using 10kDa filters, the latter resulting in the loss of many peptides and decreased levels (Supp Fig 2c).

We then confirmed the reproducibility of the guanidine HCl +80% ACN extraction method, as the %CV from 5 replicates repeated twice from different mice was around 20% for most of the peptides, with higher %CV for peptides detected with lower abundance (Supp Fig 2d), and we could also show that peptide quantification was linear with the amount of starting material in a range from 2 to 40 mg (Supp Fig 2e), indicating that tissue weight can be a satisfactory normalisation factor to compare different samples.

Tissue homogenization in guanidine HCl followed by protein precipitation with 80% acetonitrile is therefore a promising method to extract peptides from intestinal tissue for mass-spectrometry analysis, allowing the detection a large range of peptides and their robust quantification.

### Homogenization optimization

Small pieces of mouse jejunal mucosa of similar size (^~^40mg) were homogenized in different conditions in duplicates with lysing Matrix D using a FastPrep-24 for 4 × 40s at 6ms^-1^, and peptide extracted before being dried on a centrifugal concentrator. Samples were resuspended in 500μL 0.1% aqueous formic acid and purified by solid-phase extraction using a Waters HLB μElution solid-phase extraction (SPE) plate as described. Samples were analysed using high-flow rate based LC-MS method as described before.

For the acetic acid, guanidine HCl and urea extractions, tissues were directly incubated in 300μL of the lysing solution (0.25% acetic acid aqueous, guanidine HCl 6M or urea 8M) and homogenized. 200μL of the homogenate was collected into a new tube after centrifugation to remove tissue debris and lysing matrix (5min, 2000g, 4°C). 80% acetonitrile (800μL) was added to precipitate the proteins and peptides in solution were collected following centrifugation (10min, 12,000g, 4°C). For the guanidine HCl homogenates, addition of 80% acetonitrile induced a separation between an organic phase (top) and aqueous phase (bottom) in addition to the protein precipitation, with the peptides in the aqueous phase (data not shown). Peptides from guanidine HCl homogenization were also extracted using a Vivaspin 500 MCWO 10kDa (GE-healthcare) instead of using 80% ACN, collecting the flow-through after a 30 minutes spin at 16,000g at 4°C. Flow-through was then acidified adding 10% formic acid solution and extracted by SPE.

For the 80% ACN and methanol/acetic acid extraction, tissues were directly homogenized in 80% acetonitrile aqueous or 90% methanol 1% aqueous acetic acid, 200 μL of homogenate was recovered from the lysing D matrix and spun at 12,000g at 4°C for 10 minutes to separate the precipitated proteins. Peptides in supernatant were transferred to a new tube and dried before SPE extraction as previously.

For the Trizol extraction, tissue was homogenized in 500 μL Trizol and the homogenates transferred to a new tube. 100μL chloroform was added and mixed vigorously and sample centrifuged after 10min incubation at RT (15min, 12,000g, 4°C). The upper phase (aqueous) was then transferred to a new tube (from which RNA can be extracted using standard methods), 800uL 80% ACN was added to precipitate any remaining proteins and sample centrifuged (10min, 12,000g, 4°C) and supernatant dried before SPE extraction. The lower (phenol) phase was treated with 100% ethanol (1:3 ethanol:phenol phase by volume) to precipitate DNA and the mixture centrifuged (10min, 2,000g, 4°C) then transferred to a new tube. Proteins and peptides were precipitated by addition of 750 μL cold acetone and centrifugation at 7500g for 5min at 4°C. Pellets were washed once with 1mL 0.3M guanidine HCl in ethanol and air dried before being resuspended in 200uL 8M urea. Finally, proteins were precipitated with 80% ACN, centrifuged (10min, 12,000g, 4°C) and the supernatant (containing the peptides) was transferred to a fresh tube, dried and SPE extracted as previously.

---

**Figure legend (Suppl fig 3).**
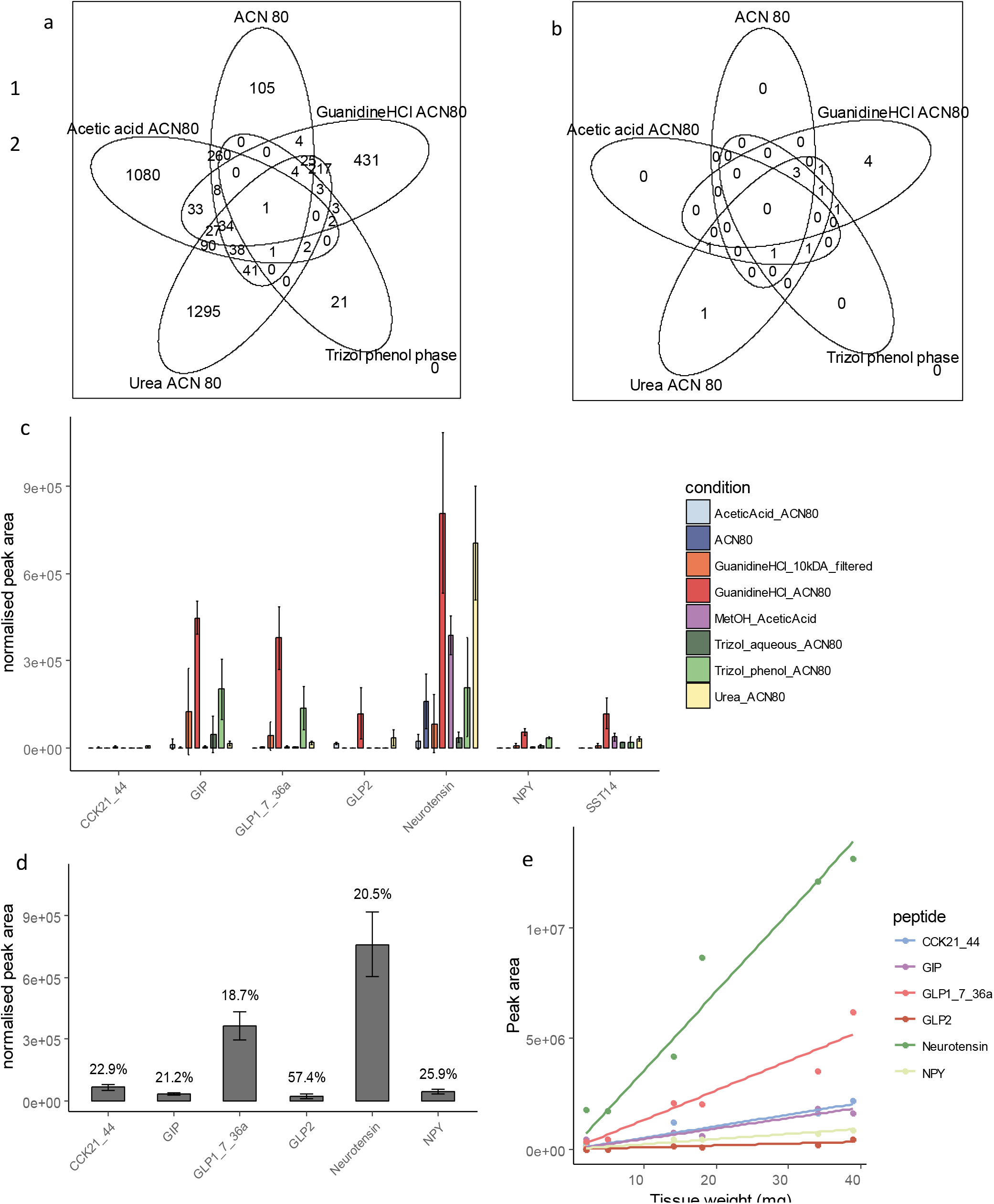
a and b: Venn diagrams showing the total number of unique peptides (a) and the number of unique peptides matching classical hormone peptides (b) found in the different methods for peptide extraction. c: Quantification of gut hormone peptides using different extraction methods, normalised by tissue weight. Data is represented as mean +/- SD from 2 replicates. d: Quantification variability for different peptides measured from 10 homogenates from 2 different mice. Data is the mean value +/- SD and values indicated above are the %CV. e: Raw peak area of peptide quantification for different samples spanning different weights from 2 to 40 mg plotted against the tissue weight. Linear regression is plotted, showing a good correlation between tissue weight and peptide quantification.

